# The landscape of the alternatively spliced transcriptome remains stable during aging across different species and tissues

**DOI:** 10.1101/541417

**Authors:** Patricia Sieber, Emanuel Barth, Manja Marz

## Abstract

Aging is characterized by a decline of cellular homeostasis over time, leading to various severe disorders and death. Alternative splicing is an important regulatory level of gene expression and thus takes on a key role in the maintenance of accurate cell and tissue function. Missplicing of certain genes has already been linked to several age-associated diseases, such as progeria, Alzheimer’s disease, Parkinson’s disease and cancer. Nevertheless, many studies focus only on transcriptional variations of single genes or the expression changes of spliceosomal genes, coding for the proteins that aggregate to the spliceosomal machinery. Little is known on the general change of present and switching isoforms in different tissues over time. In this descriptive RNA-Seq study, we report differences and commonalities of isoform usage during aging among different tissues within one species and compare changes of alternative splicing among different, evolutionarily distinct species. Although we identified a multitude of differntially spliced genes among different time points, we observed little to no general changes in the transcriptomic landscape of the investigated samples. Although there is undoubtedly considerable influence of specifically spliced genes on certain age-associated processes, this work shows that alternative splicing remains stable for the majority of genes with aging.

## Introduction

Aging is unarguably one of the most complex processes of human and animal biology and can be defined as a time-dependent, constantly weakening tissue homeostasis, resulting in various frailties and age-associated diseases. A key part in maintaining correct cellular function and thus healthiness of the organism is proper gene expression and regulation in which alternative splicing (AS) of RNA transcripts plays an important role.

AS is an important co- and post-transcriptional process in eukaryotes that enables altered forms (also referred to as isoforms) of mRNA molecules of a gene, leading to multiple mature transcripts of a single gene with possible different or modified encoded functionality^1^. Thus, AS mainly contributes to the increase of an organism’s transcriptomal diversity, allowing to synthesize multiple different proteins from a single protein-coding gene^2^. Initially, AS was considered to be a relatively rare event of genetic regulation, however, the majority of eukaryotic multi-exonic genes undergo AS^3^ and AS functions in several important biological processes, partially in a tissue-specific manner^4, 5^. The splicing process is conducted by the macromolecular spliceosome consisting of two spliceosomal complexes, the major and the minor spliceosome, which themselves are assembled from different subunits involving a multitude of proteins and maintaining RNA molecules^3, 6, 7^. Several modes of AS are known, among them exon skipping (ES), intron retention (IR), alternative 5’ splice sites (A5S), alternative 3’ splice sites (A3S), and mutually exclusive exons (MXE)1. While all of these modes may serve the same goal, *i.e.* the alteration of mRNAs by different specific mechanisms, their frequencies seem to vary in a species- and even tissue-specific way^4, 5, 8^.

Since AS provides a common level of genetic regulation in almost all biological processes, several age-related diseases are linked to either specific harmful misspliced gene variants or to a general misregulated or dysfunctional splicosomal activity. A well studied example is the Hutchison Gilford progeria syndrome, were a silent point mutation in the *lmna* gene adds a 5’ splicing signal, leading to a shortened transcript and subsequently a shorter version of its encoded protein Lamina A^9^. Because Lamina A is an important structural protein of the cell’s nucleus and its shorter version is non-functional, the nuclei of affected cells are misshaped, resulting in impaired chromatin organization and limited cell division^10^. There are many more examples of the implications of AS and age-related diseases, such as Alzheimer’s disease^11^, Parkinson’s disease^12^ and different types of cancer^13, 14^.

A few recent studies on various species and tissues investigated the change in spliceosomal activity during aging by measuring the transcription of the respective spliceosomal genes, observing an age-dependent decline in their expression^15–18^. As a common assumption, it is argued that the age-dependent decrease in the spliceosomal activity leads to an inaccurate splicing process resulting in an accumulation of non-functional mRNAs. This additional “noise” in the transcriptome might reflect another source of stress, which aging cells are exposed to, adding to the intrinsic obstacles to maintain homeostasis. Furthermore, Rodriguez *et al.* could directly report an increase in the number of alternatively spliced genes with age in different tissues of mice^19^. Nevertheless, most other studies on AS and its implications to aging are either focused on the age-dependent differential expression of splicing factors or differentially expressed isoforms of specific genes within one species and tissue^15–18, 20^. Only a small number of studies exist, that directly try to assess the general impact of aging on the heterogeneity of a cell’s transcriptome. In an RNA-Seq based study, Wood *et al.* reported changes in the usage of certain isoforms within the aging rat brain^21^. Further research is required to understand the age-related change of the isoform landscape and evaluate the true influence between AS and aging.

In the present descriptive study, we investigated and compared the global impact of the various modes of AS on the expressed transcripts in different species and tissues during aging. We analyzed a multitude of cross-sectional RNA-Seq data from four different species (*Homo sapiens*, *Mus musculus*, *Danio rerio*, *Nothobranchius furzeri*) from up to four different tissues (blood, brain, liver, skin) at four distinct ages. Whereas we identified a great number of differentially spliced genes, with many of them being involved in post-transcriptional mRNA processing pathways, we did not observe a widespread increase of spliced genes with age. Also, the average number of isoforms per gene remained constant within the investigated species and tissues. Additionally, we investigated the switch of isoforms during aging, which showed only little effect on the encoded proteins in respect to their functional domains. Finally, we could confirm a decline in the expression of major and minor spliceosomal genes and selected associated splicing factors in all investigated tissues and species, but to different extends.

We conclude that despite the acknowledged influence of single (mis)spliced isoforms on age-associated processes, AS remains in general stable and likely plays a minor role during normal physiological aging.

## Material and Methods

### Genomes and annotation

We downloaded the current genome and annotation versions of *Homo sapiens*, *Mus musculus* and *Danio rerio* from Ensembl (release 92)^22^. For *Nothobranchius furzeri*, we used the currently published assembly version of its genome and the respective annotation^23^.

### RNA-Seq libraries

All RNA-Seq libraries used are accessible at NCBI’s Gene Expression Omnibus and were published before^24^: (*Homo sapiens*: GSE75337, GSE103232; *Mus musculus*: GSE75192, GSE78130; *Danio rerio*: GSE74244 and *Nothobranchius furzeri*: GSE52462, GSE66712). For each species we used samples from up to four different tissues (blood, brain, liver, skin) at four different time points: young mature, mature, aged and old-age, see Fig. 1A for an overview.

**Figure 1.**
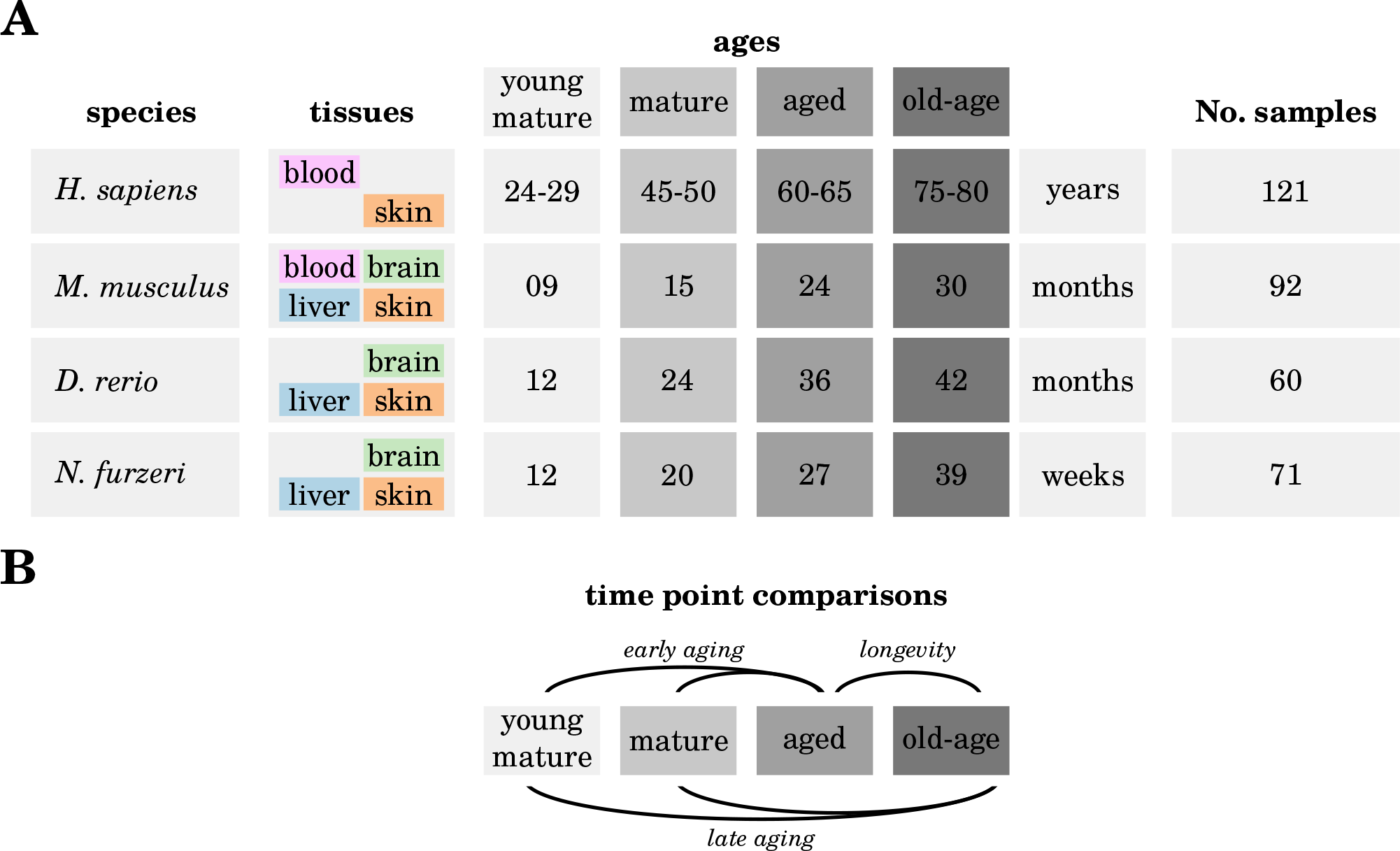
Overview of species, tissues, time points and comparisons of the utilized RNA-Seq libraries. **(A)** Samples of *Homo sapiens*, *Mus musculus*, *Danio rerio* and *Nothobranchius furzeri* were taken from up to four different tissues (blood, brain, liver, skin) at four different time points. The sampled time points can be assigned to different ages: young mature, mature, aged and old-age. **(B)** In order to explore the changes of alternative splicing, we studied three expression comparisons: early aging (young mature and mature vs. aged), late aging (young mature and mature vs. old-age), and longevity (aged vs. old-age). All comparisons were performed individually for each of the studied species and their tissues.

### RNA-Seq processing and mapping

We utilized `FastQC` (v0.11.3 ^1^) to check the read quality of all libraries before and after quality trimming, which was performed using `PRINSEQ` (v0.20.3)^25^. We trimmed reads at both sides to achieve a base quality of at least 20 and discarded all reads with a length of less than 15 nt. The quality trimmed RNA-Seq libraries were mapped onto the respective genomes using `TopHat2` (v2.1.1)^26^ with default parameters, to allow for the mapping of single reads to multiple best fitting locations and spliced reads.

### Detection of alternatively spliced genes and differentially expressed spliceosomal genes

We analyzed differential AS using `rMATS` (v3.0.8)^27, 28^. This tool has been applied successfully in other publications and showed a high precision in AS identification^27, 29^. Only results with a false discovery rate below 0.05 and an absolute inclusion level difference of AS events per sample (`IncLevelDifference`) above 0.1 were considered^29^. In addition, we manually controlled the results exemplary to ensure that AS takes place as predicted and whether it is already annotated as alternative transcript according to the respective species annotation. Furthermore, we identified the position of each AS event relative to the gene range based on the prediction of `rMATS` (see SData 1).

We selected spliceosomal genes from `Uniprot`^30^ and `Ensembl Biomart`^22^, as the macromolecular spliceosome does not only consists of proteins but also of small non-coding RNAs^5^. Based on these gene lists, we analyzed differentially expressed genes with spliceosomal activity for each species and the respective tissues. Differentially expressed genes were identified using the `DESeq2` (v1.10.0)^?^ Bioconductor package, comparing the four time points of each species and tissue, individually. False discovery rate adjustment of the resulting gene’s p-values was performed according to Benjamini *et al.*^31^. Details of all DEG results, together with the raw and normalized count values are given in detail in SData 8.

### Quantification of isoform expression

To determine the expression of all annotated isoforms independent of differential expression, we applied the tool `Stringtie` (v1.3.3b)^32^ for abundance estimation separately for each sample. We calculated only the abundance of each annotated isoform without predicting new isoforms. Normalizing the abundance of isoforms was executed using transcripts per million (TPM)^33^, we considered only those as expressed that had a TPM>1 in at least one sample. This information was used to compute the number of expressed isoforms per gene. For each gene with multiple expressed isoforms, we identified the genes predominant isoform and changes in the isoforms frequencies over time.

### Gene set enrichment analysis

Annotated gene functions of the differntially spliced genes of the early aging, late aging and longevity comparisons for each species and tissue, were obtained from the `David` functional annotation database^34^.

### Protein domain identification

For each gene that showed a switch in its predominant isoform during aging, we investigated the encoded protein domains using `Interproscan` (v5.29-68.0)^35^. For those genes with multiple predominantly expressed isoforms, we identified the corresponding protein domains and compared them among each other, counting the loss or gain of encoded protein domains of a certain gene over time.

## Results and Discussion

### Only marginally differences in the general number of expressed isoforms per gene with age

As a starting point, to investigate the general impact of aging on the heterogeneity of the transcriptome, we measured the average number of expressed isoforms per gene for each age group in all examined species and tissues (see Figure 2).

**Figure 2.**
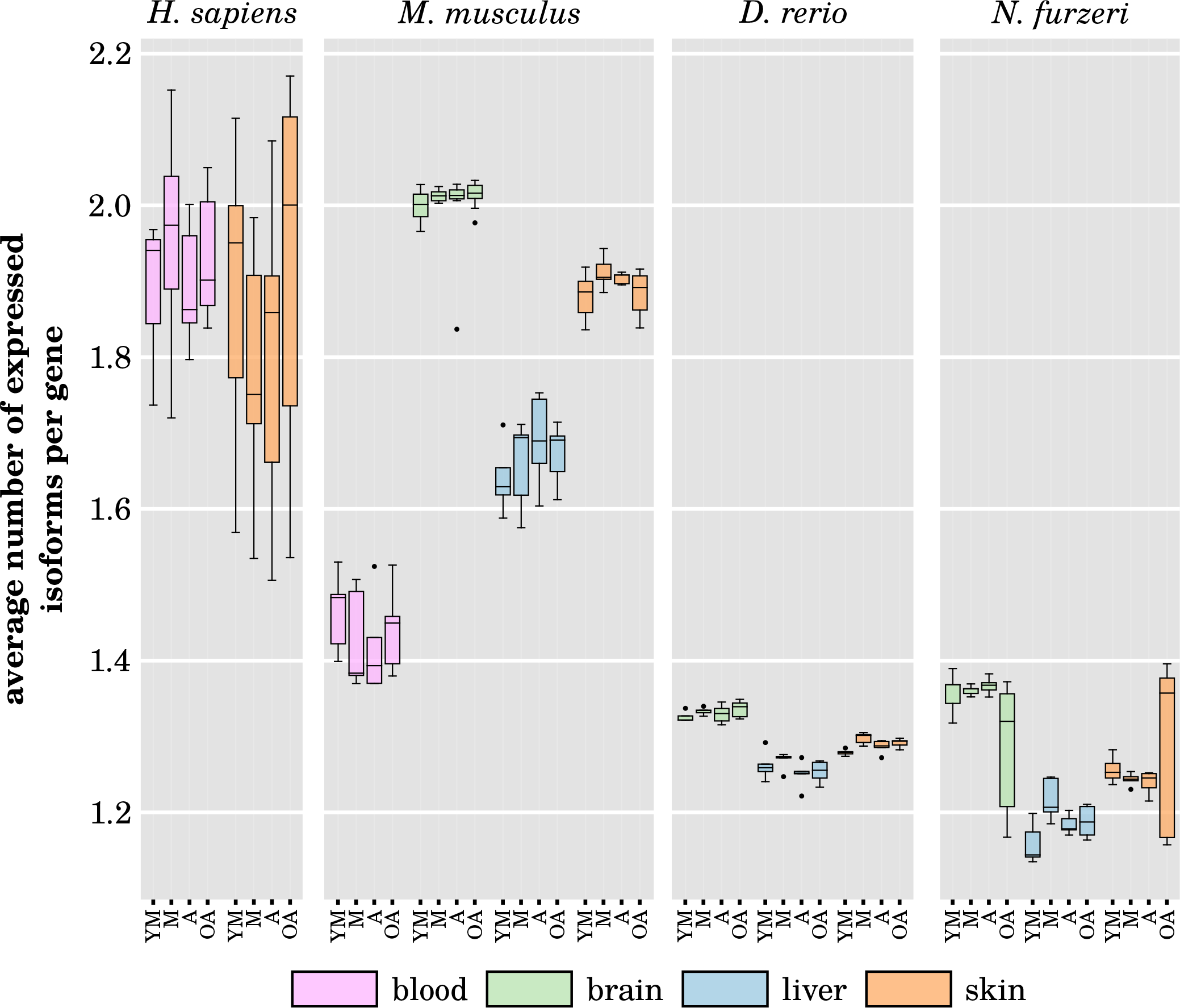
Average number of expressed isoforms per gene. The average number of expressed isoforms per gene in *Homo sapiens*, *Mus musculus*, *Danio rerio* and *Nothobranchius furzeri* for every studied tissue and time point (YM – young mature, M – mature, A – aged, OA – old-age). No significant difference in the number of expressed isoforms per gene can be observed between different ages within each investigated species and tissue. In addition, the variance between the samples of each time point and tissue remains stable, except for the old-age brain and skin sample of *Nothobranchius furzeri*, showing a manifold increased variance. The lower amount of expressed isoforms in the two fish as compared to both mammals might be biased due to the less complete isoform annotation. For more details, see SData 2 and SData 3.

Although clear differences between the tissues within each species have been observed, neither any significant age-dependent change in the average number of isoforms per gene was found, nor does the proportion of genes with multiple expressed isoforms change significantly over time (see SData 2). On average, the actively transcribed genes of both mammalians tend to express two different isoforms in parallel, while both fishes tend to have only one active isoform per gene. However, this finding could be slightly biased since the isoform annotation of protein coding genes is more comprehensive for *Homo sapiens* and *Mus musculus* than for *Danio rerio* and *Nothobranchius furzeri*, with on average about seven, five, three, and four annotated isoforms per protein coding gene, respectively. Nevertheless, there are several genes that have more than two actively transcribed isoforms during any investigated time point (see SData 3). Brain and skin samples of the *Nothobranchius furzeri* old-age time point displayed highly increased variances in the amount of expressed isoforms compared to the other time points, possibly hinting at a more deregulated splicing activity. However, this observation could not be confirmed in general. Thus, as a next step, we identified genes that either constantly increase or decrease the number of expressed isoforms with aging. Only few genes matching these criteria have been observed. *Nothobranchius furzeri* displayed the most of such genes with 14 genes showing a constant increase and 28 genes showing a constant decrease of parallel expressed isoforms (see SData 4).

Interestingly, we found many of the genes to be membrane or membrane associated proteins in all investigated species and tissues, showing this particular pattern of age-related rise/decline of isoform numbers. This is most likely due to the regulatory role of AS, which especially determines folding, topology, solubility and, thus, functions of membrane proteins, displaying its decisive task for the respective mRNAs^36^. In general, the specific annotated biological processes of the identified genes were found to be more diverse and could not be further clustered into significantly enriched molecular functions. Nevertheless, these genes still represent interesting targets for further studies, such as *foxm1* and *gas5*, which we discuss here exemplarily:

The forkhead box protein M1 that is encoded by the *foxm1* gene, represents a transcription factor of the FOX family and regulates important biological processes like cell proliferation, cell development but also play a part in longevity^37, 38^. Foxm1 has a key role in cell cycle progression as well as chromosomal segregation and genomic stability and therefore is tightly linked to the age-associated processes of cellular senescence and cancer formation^39, 40^. Three different isoforms are known to be transcribed from *foxm1*, where one acts as a transcriptional repressor (isoform A) and two as activators (isoform B and C), associating the latter two to the proto-oncogenic character of *foxm1*^40^. We observed a constant age-dependent decrease in the number of expressed isoforms of *foxm1* in the mice skin samples, with all three isoforms being actively transcribed in the first two mature time points then being reduced to two in aged mice and finally just one in the old-age time point. When neglecting the TPM low expression threshold, the same trend could be observed for the human skin samples. All other tissues showed a constant but relatively low expression of just one isoform within all age groups, with exception to the skin samples of *Danio rerio*, expressing the opposite pattern during aging, with just one actively transcribed isoform at the young mature time point and rising to have all three isoforms expressed at the latest old-age time point. This observation might suggest an age-related control of splicing of *foxm1*. For *Nothobranchius furzeri*, however, only one known isoform is currently annotated, so no predication could be made.

The other interesting gene, *gas5*, does not encode a protein but a long non-coding RNA (lncRNA), which was shown to be involved in apoptotic and tumor formation processes^41^. Several different isoforms are known and predicted for *gas5*, of which some host various small nucleolar RNA (snoRNA) sequences^42, 43^. But expression of *gas5* is also strongly linked to growth control and senescence of human T-cells, indicating a potential role in the age-associated phenomenon of immunosenescence^44^. We observed a similar age-dependent decline in the number of transcribed isoforms of *gas5* in the liver and skin samples of *Mus musculus* and *Danio rerio*, showing the lowest number of parallel actively expressed isoforms at the old-age time point. On the opposite, the brain samples of both species and both investigated human tissues show a more diverse pattern in the amount of expressed distinct isoforms, but always with the highest number during old-age. The exact function and role of the different isoforms of this lncRNA during aging remains to be explored, but it serves as an interesting potential target of further research. Again, no homologous gene is currently annotated or known for *Nothobranchius furzeri*.

### Changes in the expression of the main isoforms do not disrupt the encoded functional protein domains

Due to the fact that the number of different expressed isoforms and genes do not change significantly with age, we investigated if the mainly expressed isoforms of all genes are switched in an observable age-related fashion (see Fig. 3). For this analysis, only single isoforms with a higher expression than other expressed isoforms were taken into account^45^.

**Figure 3.**
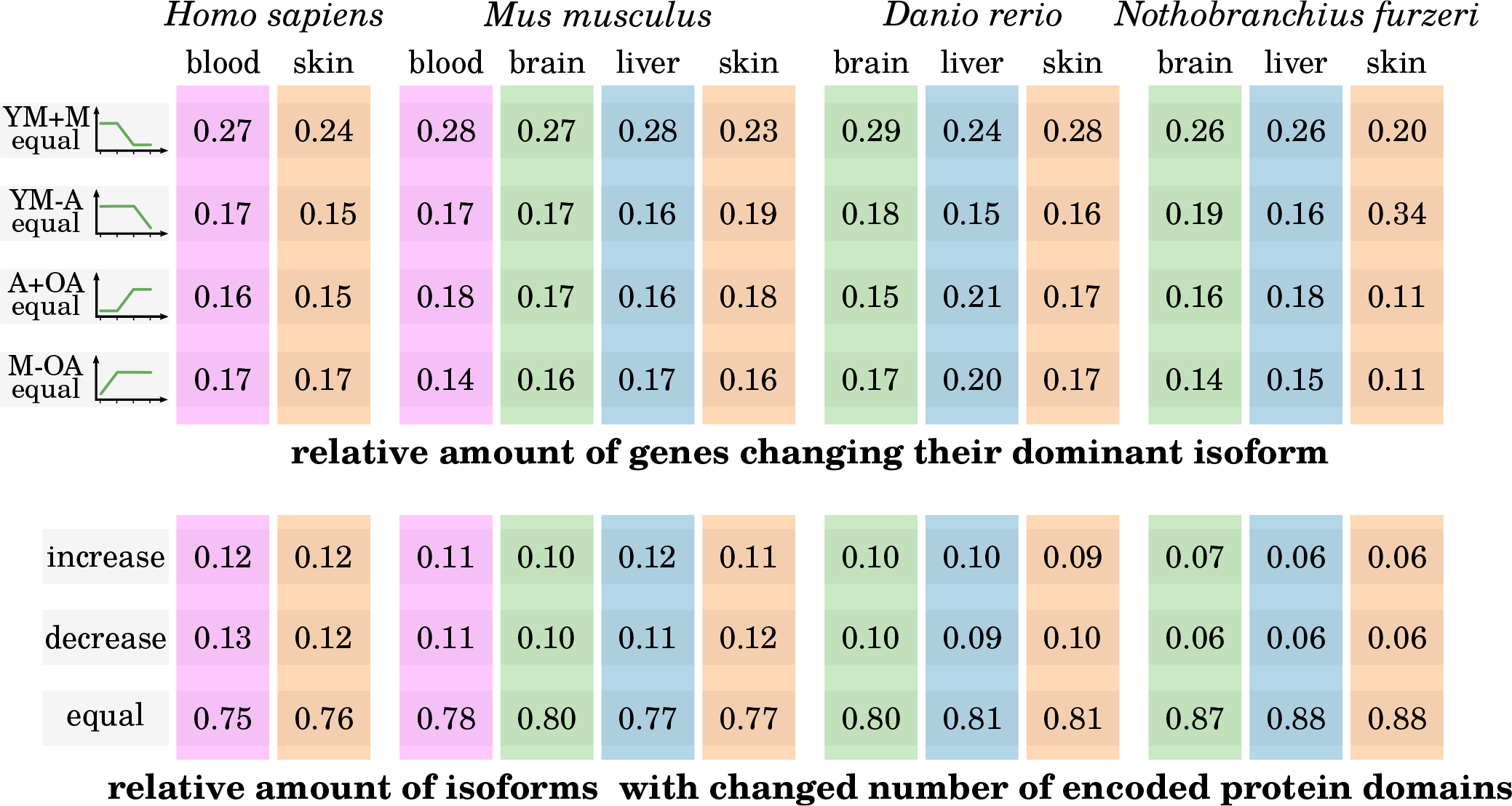
Frequency of isoform switches during aging and impact on the amount of encoded functional domains. **Top:** The relative amounts of genes for each species and tissue that switch their main isoform in one of four age-dependent manners are displayed. The dominant isoform switch occurs after the mature time point (YM+M equal), it remains the same until the aged time point and switches during the old-age (YM – A equal), it remains stable from the aged to the old-age time point (A+OA equal), or it switches directly after the the young mature time point (M – OA equal). Generally, age-dependent isoform switching occurs only for a minority genes and takes place more frequently in later time points. **Bottom:** The relative amounts of isoforms, which encode less, more or still the same number of functional protein domains after being switched with age. Only genes with a changing dominant isoform are considered.

We observed an aging-dependent switch in the dominantly transcribed isoform for 11 – 34 % of all expressed genes in all investigated species and tissues. Interestingly, with about 26 %, the most of all expressed genes switched their main isoform at the aged time point after being unchanged during both mature time points, regardless of tissue and species, see Figure **??**. The same species- and tissue-independent relative amounts can be observed for genes that switch their main isoform directly after the young mature time point or at the latest old-age time point, with around 16 – 18 % of the relevant genes for each of these cases individually. We examined these genes with altered predominant isoforms during aging more closely, by determining the amount of encoded functional protein domains in the different dominant isoforms. Functional protein domains (in short functional domains or protein domains) are specific units of amino acid sequences, which are structurally and functionally conserved and determine the precise function of proteins^46^. Surprisingly, in around 80 % of all genes showing an age-dependent isoform switch, the encoded number of protein domains remained unaltered, with the remaining genes showing equally fractions of isoforms with increased or decreased encoded functional domains after being switched (see Figure 3). This again holds for all examined species and tissues. A change in the amount or composition of encoded protein domains within an mRNA has great influence on the topology, localization and function of the translated protein. However, since only a small fraction of switched mRNA isoforms is concerned by such a change in an age-dependent manner, most AS events seem to affect either sites within the mRNAs that encode unstructured protein regions or untranslated regions (UTRs). In fact, it was recently reported that changes in trailing 5’ and 3’ UTRs as well as intrinsic unstructured regions of mRNAs due to AS are over-represented as compared to changes altering encoded protein domains^46^. Still, changes within the 5’ or 3’ UTRs can have significant effect on the translation of the respective mRNA, because the primary sequence and secondary structure of UTRs regulate the transcript’s stability, turnover and efficiency of translation^47^. Yet, since most of the age-dependent isoform switches are subject to non-obvious functional changes, they might influence the effective translational availability rather than their abundance, the real impact of aging on the process of AS is hard to assess.

Nevertheless, for individual genes, the change in the switched isoform can be more drastic with respect to their encoded function and the proportion of different expressed isoforms of a single gene could have significant impact on several aging-related processes^48^. For example, we found the mainly expressed isoform of the *bcl2l1* gene to switch at the aged time point of the mouse liver samples from encoding is short protein version (Bcl-xS) instead of its long version (Bcl-xL). Whereas Bcl-xL is an apoptosis inhibitor, the short version Bcl-xS is an apoptosis activator, displaying the AS controlled age-dependent transition from repressing to activating apoptotic processes in hepatic cells. Apoptosis is an important mechanism in aging liver tissue to maintain homeostasis and several dysfunctions, and diseases of the liver are related to programmed cell death processes^49, 50^.

Also, amongst others, *srsf1* and *srsf6* change their predominantly expressed isoform over time in different murine tissues, and *srsf3* in human skin. These genes encode for SR proteins that are known to regulate certain age-related genes^51^. For example, SRSF1 and SRSF3 have direct impact on the expression of the important tumor protein p53 that regulates the cell cycle, senescence and apoptosis^48, 51^. In addition, the gene *gpr18* in human blood has been identified to change its isoform ratio^15^, which is confirmed by our data. Although the isoforms of these genes are not differentially alternatively expressed, the switched expression of their mainly transcribed isoforms can influence the regulation of aging and senescence.

### Less than 5 % of transcribed genes are differentially alternatively spliced during aging

To further assess the influence of aging on the splicing process, we investigated the expression change of differentially alternatively spliced genes (DSGs) between three different comparisons: early aging, late aging and longevity (see Figure 1). For every species and tissue, we observed quite varying amounts of DSGs and in contrast to previous findings^19^, we could not confirm a general increase of DSGs with age (see Figure 4A). Rodriguez *et al.* compared the number of DSGs of murine tissues, but between different ages (4 months to 18 months and 4 months to 28 months) as compared to our time points (9/15 months to 24 months and 9/15 months to 30 months), which could explain this discrepancy. In addition, they investigated different tissues (skeletal muscle, thymus, adipose tissue and bone, besides skin tissue) and analyzed microarray data instead of RNA-Seq data, adding to the factors that make a direct comparison difficult. The only similar observation they made was an increase of DSGs in their late aging comparison (4 months to 28 months) as compared to their early aging comparison (4 months to 18 months). In general, within our data we observed only few DSGs in the mammalian samples (less than 2 % of the transcribed genes) compared with the fishes samples (less than 5 % of the transcribed genes) and highest number of DSGs within the skin tissues, with exception to *Danio rerio*, having more age-dependent spliced genes in brain (311 DSGs) than skin (241 DSGs). This might indicate that the tissue-specific regulation of AS processes prevails over any possible common aging effect. When analyzing differential expression of splicing within our age comparisons, we also investigated the occurring AS modes (exon skipping (ES), intron retention (IR), alternative 5’ splice sites (A5S), alternative 3’ splice sites (A3S), and mutually exclusive exons (MXE)) that contributed to the identified DSGs (see Figure 4B). We identified ES as the most prevalent pattern in all species, which is in agreement with previous findings^1^. There are only a few exceptions where one of the other AS modes prevails (see SData 5): In the human blood samples, IR is the main form of the alternatively spliced genes during early aging. Interestingly, A5S and A3S make up more than half of the AS modes in the longevity comparison in human. In the mouse blood samples, a decline from 86 % ES splice events in the early aging comparison to only 20 % in the longevity comparison can be observed, compensated by a rise of MXE splice events to 60 %. The same shift from mainly ES to MXE splice events can be observed in the murine skin samples. In contrast to any of the other AS modes, MXE is a relative complex and controlled splicing process and less likely to happen accidentally due to a deregulated splicing machinery^52^, suggesting a stable working spliceosome even with high age. In *Danio rerio*, ES remains the main AS mode within all age comparisons and investigated tissues with a share of 54 % – 79 % of all occurring splice events. The fraction of AS modes does not change considerably in the brain and skin of *Nothobranchius furzeri*, with the notable exception of A5S and A3S, which seem to occur more often during early aging than late aging in the brain samples. Notably, the amount of IR increases during aging, especially in the liver samples of *Nothobranchius furzeri*, although this AS mode is known to be rare in vertebrates^1^. In general, the occurrence of the different AS modes seems to be tissue-specific with no observable distinct drift of specific splicing events during aging, let alone a common tendency between the individual tissues of one of the investigated species. In addition, between the four species, there are only few commonalities, besides ES being the predominant splicing event and the liver samples showing an unexpectedly high rate of DSGs derived from IR splice events. However, given an age-dependent decline in the correct functionality of the splicesome, one could expect a systematic decrease of more controlled splicing events, such as MXE.

**Figure 4.**
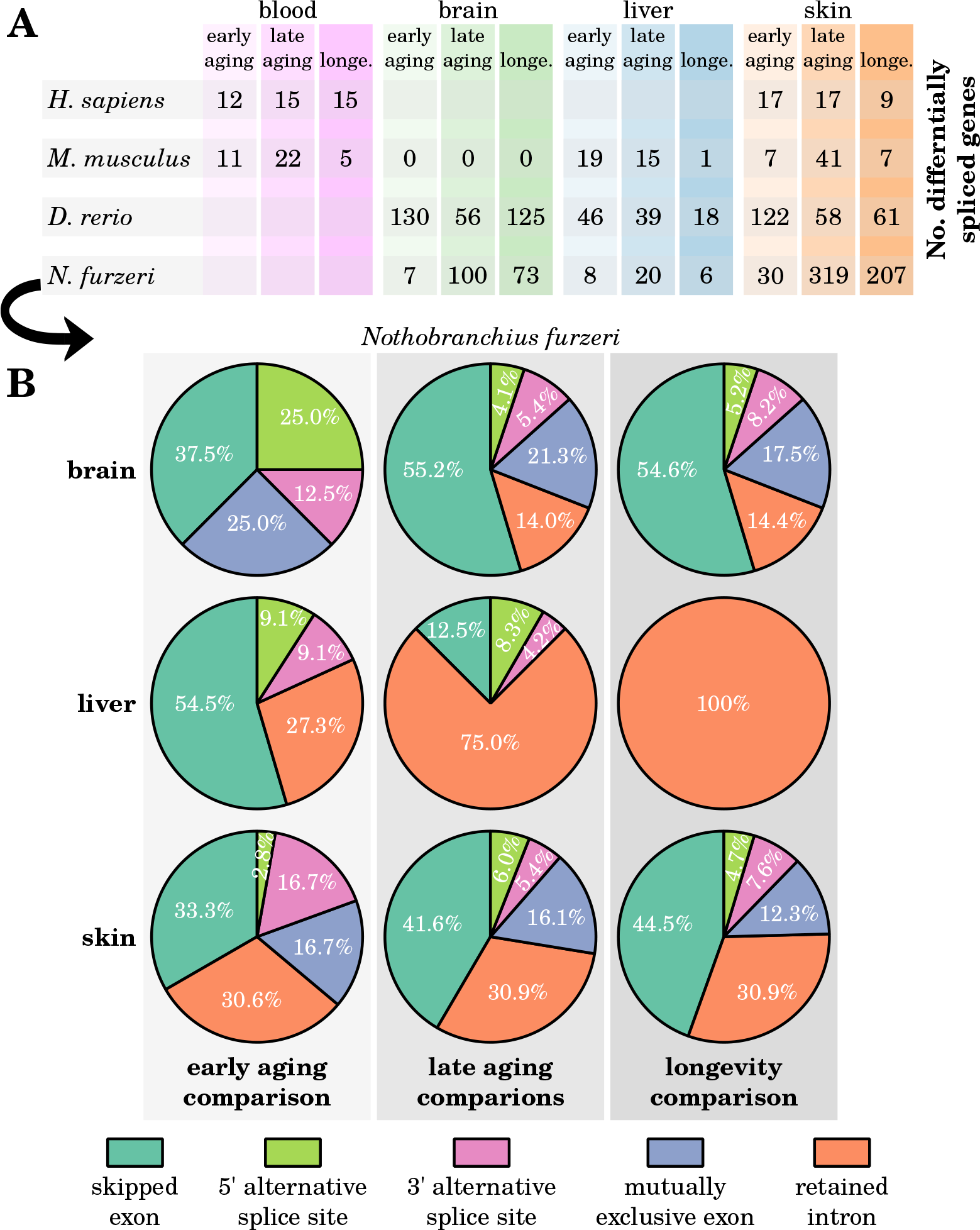
Number of identified differentially spliced genes. **(A)** The amount of differentially spliced genes for the early aging, late aging and longevity (longe.) comparisons for every investigated species and tissue are given. Far less differentially spliced genes could be observed for *Homo sapiens* and *Mus musculus* as compared to the fishes *Danio rerio* and *Nothobranchius furzeri*. Also, no general trend of increased gene splicing can be recognized in any of the species or tissues. On the arbitrary example of *Nothobranchius furzeri*, **(B)** depicts the fractions of the exact AS modes (skipped exon, 5’/3’ alternative splice site, mutually exclusive exon, retained intron) that contribute to the identified differentially spliced genes. Despite skipped exon being the most abundant AS mode in almost every age comparison, each tissues displays its own signature of splicing events during aging. More detailed information, including the AS modes of the other species can be found in SData 5.

Furthermore, we observed that AS tends to either take place in the middle or towards the 3’ end of the expressed transcripts, regardless of the splice event, in all investigated tissues and species. In *Nothobranchius furzeri*, even the majority of AS events take place close to the 3’ ends of the spliced transcripts (within the last quarter of the transcripts sequences), whereas for the other three species most AS events occurred in the middle of the spliced RNAs (within the second and third quarter of the respective sequences). This is not completely unexpected, since AS mostly changes protein characteristics only slightly1, 5. Changes in the primary sequences of transcripts near the 5’ end due to AS, resulting in a frameshift, have a higher chance to alter the encoded protein function more significantly, which is commonly believed to be avoided. Again, if there is a general failure of the splicing machinery with increasing age, we would anticipate to see AS events to occur more uniformly distributed in transcribed sequences.

### Age-related alternatively spliced genes are tissue- and species-specific

When comparing the observed DSGs between the tissues for each species individually, we found almost no overlap between the tissues (see Figure 5 and SData 6), confirming earlier findings of age-related tissue-specific AS regulation^1, 19^. Nevertheless, a few common genes could be identified being altered by AS during aging. For example, during early aging in both the liver and skin of *Mus musculus* the ubiquitously expressed gene *hnrnpa2b1* undergoes AS. It encodes for a member of the heterogeneous nuclear ribonucleoprotein family. Besides being involved in many different cellular functionalities, such as stabilization of telomeres, cell proliferation and splicing of other pre-mRNAs, it is associated to several diseases^26, 53^. *Hnrnpa2b1* has several known isoforms with most of them coding for the same protein, but one non-functional transcript (ENSMUST00000204090) is expressed highly in both tissues. Whether alterations in this genes isoform expression have any influence on further age-related AS remains an interesting open question. Furthermore, it seems to have at least some connection to aging because it was associated with longevity in a previous study^16^.

**Figure 5.**
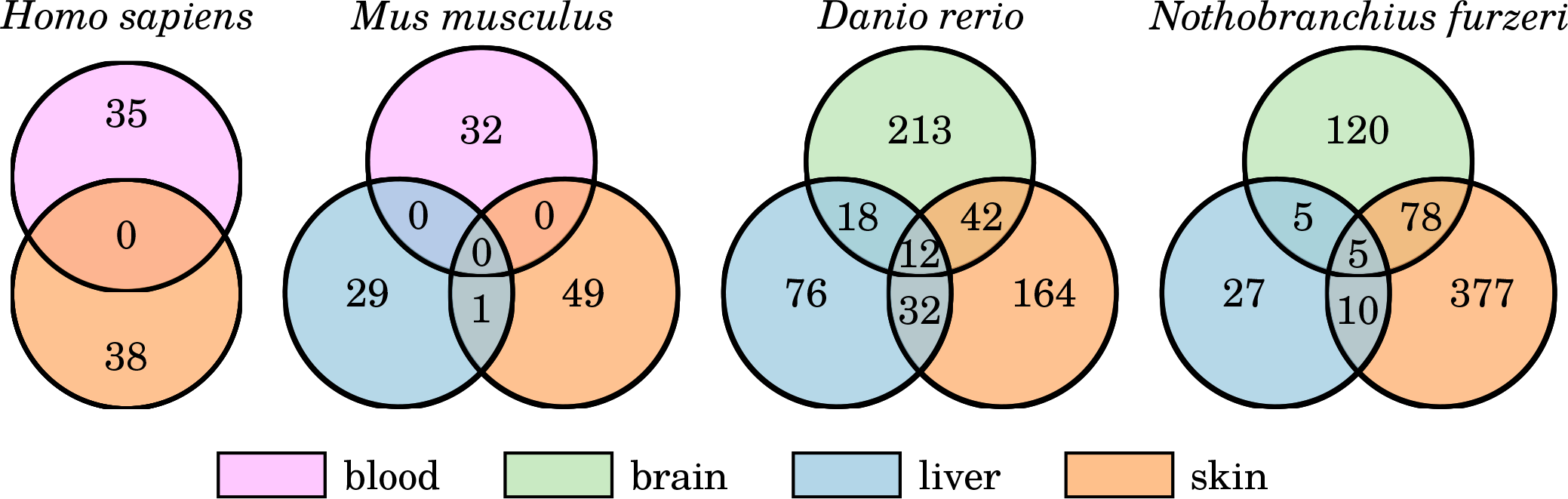
Overlap of DSGs between the different tissues within each investigated species. The comparison of the age-related DSGs demonstrates the tissue-specificity of the AS processes, showing only little overlap within each species. Note that the different age comparisons of each tissue are collapsed. For further results, see SData 6

But also many commonly shared DSGs in both fish species have functions in the splicing process or mRNA processing, like *u2af2b*, occurring to be alternatively spliced in all *Danio rerio* investigated tissues during early aging, encoding for an important subunit of the spliceosome^54^.

Even fewer overlap of DSGs can be found among the four different species (see SData 6). The only shared alternatively spliced gene in the aging human and mouse blood samples was *rpl13a*, encoding for a ribosomal protein, which was reported to play a special role in specific translational control of inflammation associated genes, next to its normal function as part of the ribosome^55^. This is of particular interest as inflammation processes tend to be deregulated with rising age in vertebrates and being implicated in many age-associated disorders^56, 57^.

Most of the few shared DSGs in the skin samples of *Danio rerio* and *Nothobranchius furzeri* were observed to be cell structural genes. However, one more interesting gene, *tp63*, was found to express different isoforms in both fish species during aging. It encodes for a protein named Tumor protein 63 (or transformation-related protein 63), which is a homolog of the intensively studied tumor suppressor gene p53^58^. There are two distinct major isoforms, TAp63 and *δ* Np63, with the first regulating apoptotic processes^59^ and latter being involved in skin development and stem cell regulation^60^. Interestingly, we see a switch to the TAp63 isoform towards the old-aged individuals. Since it was recently shown that TAp63 prevents premature skin aging by supporting the maintenance of adult stem cells, age-related AS of this gene might have a conserved role in longevity^61^.

### Alternatively spliced genes heavily function in mRNA processing and transcription regulation besides tissue-specific processes during aging

To infer the potential role of the identified aging-dependent DSGs, we systematically examined their annotated functions for the individual tissues and species (see SData 7). Most interestingly, we found that genes directly responsible for splicing, post-transcriptional mRNA processing, transcriptional and translational regulation are differentially spliced in almost all aging comparisons (see Figure 6). Most of these DSGs include members of the DEAD box protein family, ribosomal proteins, eukaryotic initiation/elongation factors or splicing factors of the SR family. Similar observations were already made in different mouse tissues and human blood samples^15, 19, 51, 62^. In addition to our observations, it shows that age-related AS of various splicing and transcription components is, to some extend, a conserved aging feature. Since we did not observe a drastic change in the heterogeneity of expressed isoforms with age, we assume that the alterations of these splicing and transcription components do not lead to a generally disturbed spliceosomal or mRNA processing activity. Transcription and splicing are two closely connected processes, wherein several of the splicing factors directly interact with the transcription machinery^63^. Potentially, AS induced age-related changes in some components of the spliceosome might reflect adaptation mechanism to an altered transcription process. However, to understand the consequences of these changes, further research is required on the exact effects for the encoded proteins of the respective DSGs.

**Figure 6.**
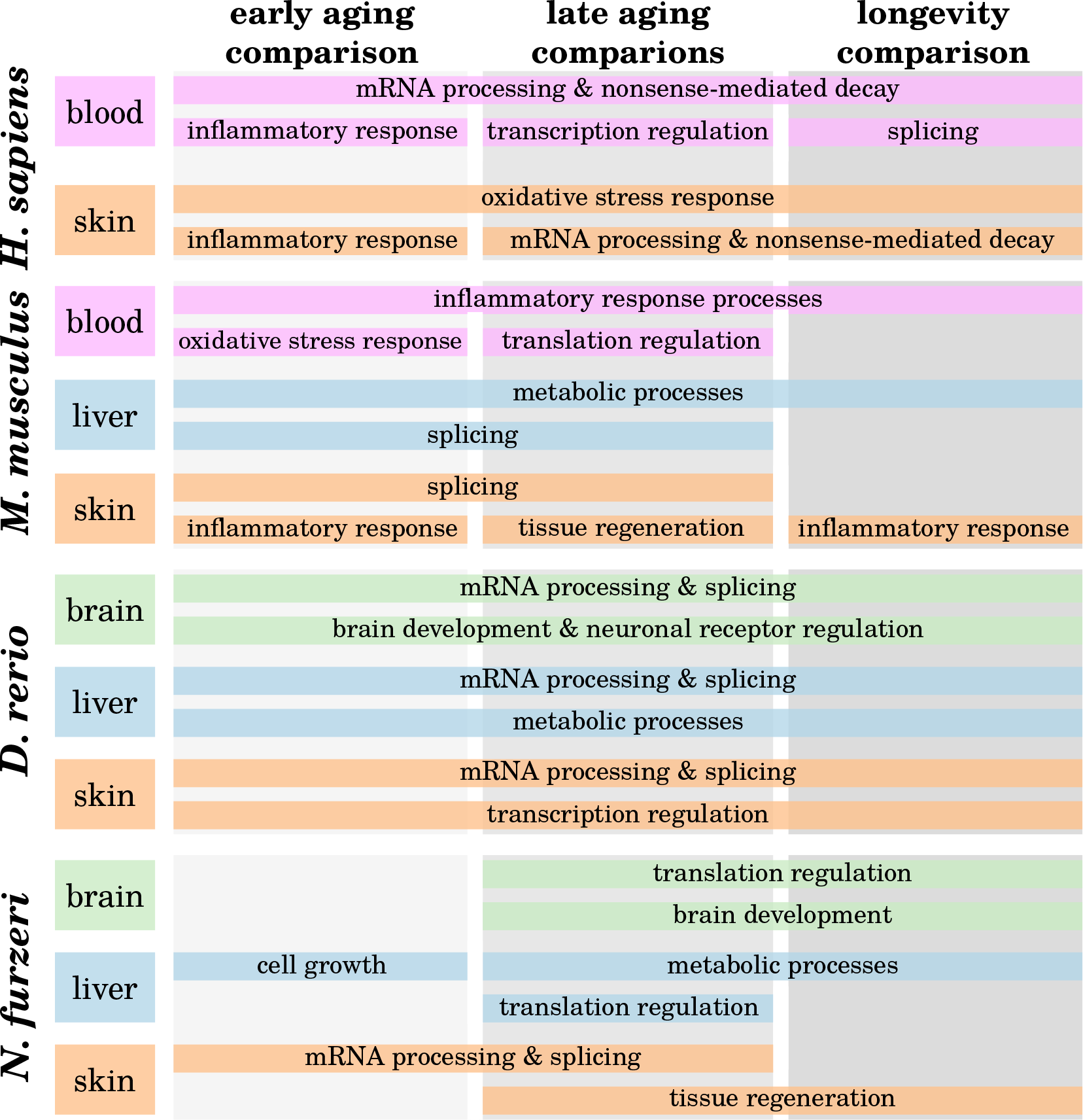
The prevailing biological functions associated with the alternatively spliced genes in the examined age comparisons. Interestingly, various DSGs involved in splicing, post-transcriptional mRNA processing as well as transcriptional and translation regulation were observed in almost all aging comparisons, regardless of the investigated tissue and species. Among others, tissue-specific genes functioning in aging-associated processes were found to be subject to AS, such as tissue regeneration, inflammation or oxidative stress response. Detailed information can be found in SData 7.

In the human blood and skin samples, we additionally observed many age-related DSGs to be involved in the process of nonsense-mediated mRNA decay (NMD). NMD, which is the degradation of mRNA transcripts due to premature termination signals^64^, was itself observed to be regulated by AS through aging in the human blood and skin. Whereas the main function of NMD is to discard aberrant mRNAs that encode potentially deleterious proteins, it is also used as a transcriptional regulatory level, controlling stability and availability of mRNAs by AS^65^. Possible implications of an altered NMD mechanism through age-related AS remain again an interesting target of further studies.

Besides these common and partially conserved findings, we observed also many DSGs to function in tissue-specific aging-associated processes. AS of genes involved in inflammatory response were identified mainly in the human and mouse blood and skin samples, most likely being associated to the age-related increase of inflammatory processes in these tissues^56, 57, 66^, since AS is extensively used to increase the isoform diversity of genes of the immune system^67^.

As expected, the liver samples of mouse and both fishes display DSGs of various metabolic processes, hinting at possible alterations in digestion or detoxification with age. In the brain samples of *Danio rerio* and *Nothobranchius furzeri*, many DSGs are associated with neuronal receptor regulation and brain development processes. It is a known feature of teleosts, such as both investigated fishes, to maintain neurogenesis even in older ages^68, 69^.

### Age-related changes in spliceosomal activity correlates with the number of differentially spliced genes

The spliceosome is a ribonucleoprotein complex, consisting of a high number of components to ensure the correct process of splicing with a high accuracy including several control mechanisms and different modes^3, 5^. As a further step to understand how AS could be affected by aging, we analyzed the expression of the main protein-coding and non-coding RNA genes of both the major and minor spliceosome as well as transcriptional changes of associated splicing factors (see Figure 7 and SData 8). The deregulation of the spliceosome or at least some of its components can have severe consequences for the organism and was focus of recent aging-related microarray-based studies^16, 17, 51^.

**Figure 7.**
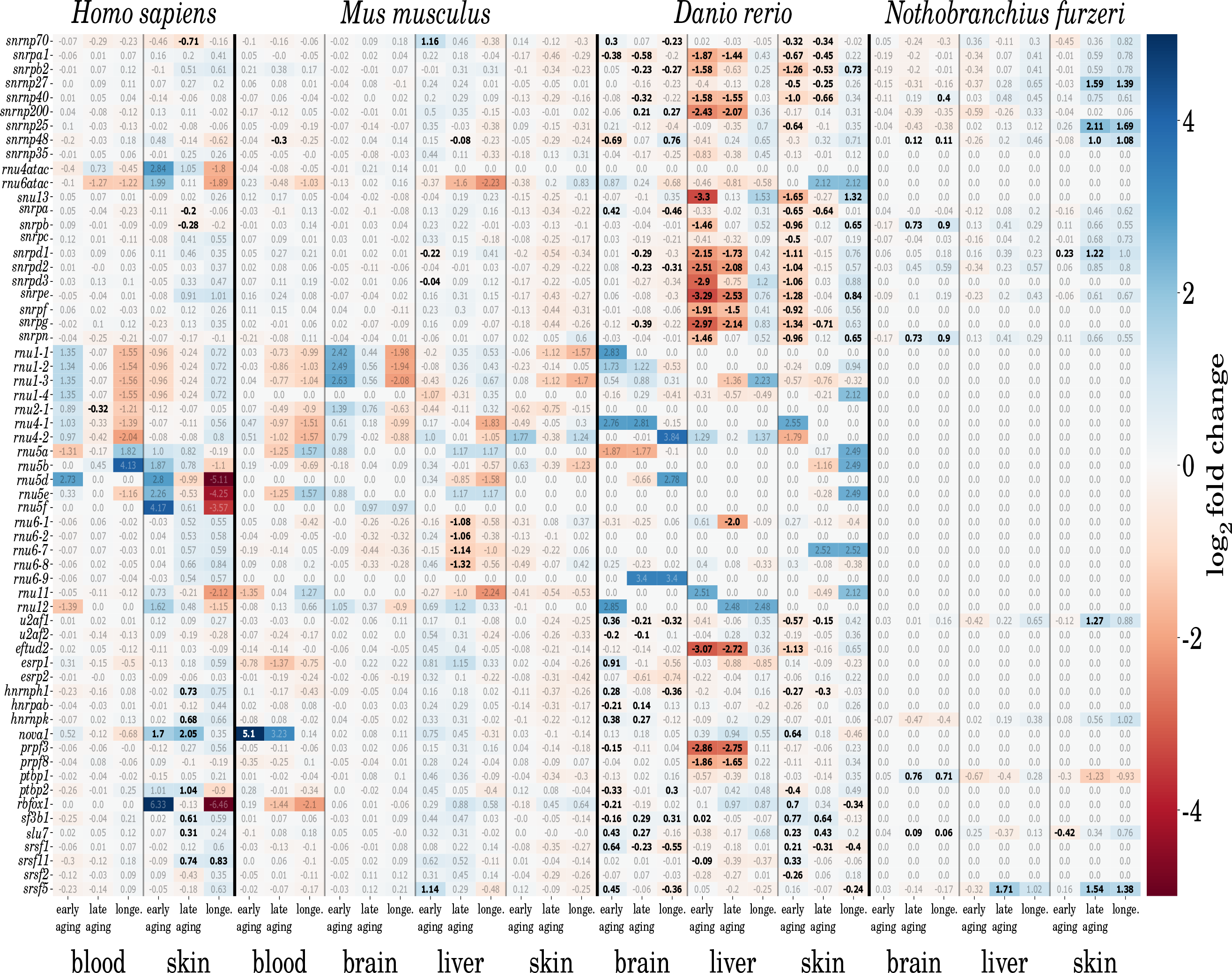
Transcriptional changes of the major and minor spliceosomal genes as well as splicing factor genes during aging. Numbers indicate *log2* fold changes between two compared ages, were a positive value indicates an upregulation and a negative value a downregulation of the respective gene with aging. Bold numbers mark statistically significant changes.

For human and mouse, we observed a stable expression of the main spliceosomal genes during aging with only a few significant changes. In human blood, only the U2 small nuclear RNA is significantly downregulated towards the old-age time point. Other RNAs, such as U1 and U4, show the same tendency. In skin, the gene *snrnp70*, encoding for an important subunit of the small nuclear ribonucleoprotein U1, is down-regulated in the late age comparison. Interestingly, the same gene shows an about two-fold upregulation in the mouse liver within the early aging comparison, whereas the small nuclear RNA U6 becomes downregulated in the long-lived individuals. All of the mentioned genes are crucial for the correct assembly and function of the major spliceosome, but their change during aging seems to be little in both mammals. Still, several additional splicing factors show a strong upregulation with age in the human skin samples, such as *nova1*, which is also highly upregulated during early aging in the mouse blood samples. The equally named encoded protein of *nova1* was first thought to be a brain-specific splicing factor^70^, but was later discovered to play a key role in the splicing of pancreatic beta cells, too^71^. Here, we observe that it also seems to be active in two more tissues, human skin and murine blood, in an aging-related manner.

However, both investigated fishes present a different picture. Whereas *Nothobranchius furzeri* shows as few significant changes as both mammals, important genes of the major and minor spliceosome are rather up-instead of downregulated during aging. In its skin, we observed an upregulation of the gene *snrnp27*, which is part of the major spliceosomal U4/U6 subunit, and the two genes *snrnp25* and *snrnp48*, which are part of the minor spliceosomal-specific U11/U12 subunit. Additionally, the two spliceosomal associated genes, coding for the ribonucleoproteins SNU13 and SNRPD1 are also more than two-fold increased in their expression. All of these genes show a significant higher activity towards the old-age time point and may indicate a need for an enduring splicing activity. Also, in the brain of *Nothobranchius furzeri*, a significant activation of spliceosomal related genes, like *snrnp40* (ribonucleoprotein U5) or *snrnp48* (ribonucleoprotein U11/U12) was observed with age. However, the transcriptional changes were not as strong as compared to the skin.

In contrast to the other examined species, *Danio rerio* displays extensive changes in the expression of its spliceosomal genes and associated splicing factors during aging. It displays a distinctive down-regulation of most of the ribonucleoprotein coding genes with age. This can especially be seen in the liver and skin samples during the early and late aging comparison. Most interestingly, the longevity comparison between the aged and the old-age animals revealed either no change in expression or a slight upregulation of some ribonucleoproteins, indicating no or only minor further modulations of the spliceosome after a certain age is reached. The brain shows in principal the same pattern as the other tissues, but similar to *Nothobranchius furzeri* to a much weaker extend. If this strong downregulation can be correlated to the comparatively high amount of significantly spliced genes in *Danio rerio*, remains an interesting open question.

In general, we do not identify a significant increase or decrease of expression of the catalytic units of the minor spliceosome (U11, U12, U4atac, U6atac) during aging.

## Conclusion

Within this descriptive RNA-Seq study, we analyzed the impact of aging on AS in four evolutionarily distinct species and up to four different tissues. For each species, we investigated samples with five or more replicates of two mature (one young and one middle-aged), an aged and an old-age time point. When comparing these ages among each other, we observed a varying amount of significantly differntially spliced genes (DSGs) for the four species and tissues. Whereas the fishes *Danio rerio* and *Nothobranchius furzeri* showed in total several hundred DSGs, much less were identified in *Homo sapiens* and *Mus musculus*. However, a general increase of AS with age could not be observed, which is in contrast to earlier reports^19^. Most likely, these differing results arise from the different experimental setups. The study of Rodriguez *et al.*^19^ was based on microarray data of only murine tissues (skeletal muscle, thymus, adipose tissue, bone and skin) and completely different time points (4 instead of 9 months and 18 instead of 24 months for the mature and aged time points respectively) as compared to our data. They observed an increase of AS in the bone, muscle and skin tissues with age, which we could confirm at least for the latter tissue, but to a much lower extend. Taking other tissues and species into consideration, their observation of an age-related accumulation of DSGs cannot be generalized.

Interestingly, even though we identified age-related DSGs to be almost exclusively tissue- and species-specific, many of them function in various post-translation mRNA processing steps, transcriptional and translational control and splicing processes themselves, independent of the investigated species and tissues. This observation indicates AS and other RNA processing mechanisms to be indeed modified with normal physiological aging, and that this feature seems to be conserved to some extend. Other studies observed similar results in human and mice^15, 19, 51, 62^. As a consequence, the specificity of the spliceosome and thus transcription of specific gene isoforms seem to change during aging. Besides mRNA regulatory processes, age-related DSGs are mainly involved in tissue-specific aging-associated processes, such as inflammation in the mammalian blood and skin samples^56, 57^, metabolic process in liver and tissue regeneration processes in the fishes brain and skin samples^68, 69^. However, since more aging-related genes are expressed with increasing age, it is not totally unexpected to observe these genes to be processed by AS.

Other recent microarray-based studies focused on the expression changes of spliceosomal genes and splicing factors, reporting a decline in the expression of several components of the splicing machinery^16, 17, 51^, possibly resulting in an increased heterogeneity of isoforms and transcriptional noise^72, 73^. When examining the expression activity of major and minor spliceosomal genes and several splicing factors, we observed only a weak downregulation of these genes during aging in any of the investigated human and mouse samples and even a slight upregulation in *Nothobranchius furzeri*. In contrast, *Danio rerio* displays massive transcriptional modulation of spliceosomal genes and associated splicing factors during aging.

One of the most important findings of our study is that these age-related changes in the splicing process, either by self-regulation through significant changes in the expressed isoforms of various spliceosome components or the up-/downregulation of spliceosomal genes are not reflected in the general landscape of present RNA transcripts. Neither does the number of expressed isoforms per gene change significantly over time, nor are the proteins encoded by genes that switch their predominant isoform during aging affected. Nevertheless, different isoforms expressed by the same gene can still encode exactly the same protein and changes in their untranslated regions can have major impact on their stability and translation efficiency^46^. Further studies based on single cell RNA-sequencing could give deeper insights in the diversity of the transcriptome during aging and its implication in biological processes^74^.

In conclusion, while the overall landscape of the aging transcriptome seems to be unaffected by aging-related changes in AS or the spliceosomal activity, single alternatively spliced genes can have crucial influences on certain aging processes but AS shows not to be deregulated in general with increasing age.

## Acknowledgments

We thank I. Goerlich and I. Heinze for their skillful and diligent technical assistance in NGS data acquisition and M. Groth and M. Platzer for providing RNA-sequencing datasets. We thank D. Esser and P.A. Irizar for their help in method evaluation, and all participants of the HACKEN 2017 project.

## Author Contributions

*Conceptualization:* EB, PS, MM. *Data curation:* EB, PS. *Formal analysis:* EB, PS. *Data interpretation:* EB, PS. *Methodology:* EB, PS. *Resources:* MM. *Validation:* EB, PS. *Visualization:* EB, PS. *Writing – Original Draft Preparation:* EB, PS. *Writing – Review Editing:* EB, PS, MM. *Project Administration:* MM. *Supervision:* MM. *Funding acquisition:* MM.

## Additional information

All relevant data can be found within the paper and its supporting information files. The online supplement is integrated into the open science framework (https://osf.io/) and and can be found at https://osf.io/rz6kc/ or its corresponding DOI 10.17605/OSF.IO/RZ6KC.

## Footnotes

1 www.bioinformatics.bbsrc.ac.uk/projects/fastqc

